# Mitochondrial genomes of individual microfilariae: Undescribed *Dirofilaria*-like filariae from Malaysian cats and two filaria species related to *Dirofilaria* and *Mansonella* from Indonesian macaques

**DOI:** 10.64898/2026.02.09.704775

**Authors:** Irina Diekmann, Young-Jun Choi, Taniawati Supali, Rahmat Alfian, Yossi Destani, Elisa Iskandar, Noviani Sugianto, Mohd Hatta Abdul Mutalip, Nor Azlina Abdul Aziz, Khairiah Ibrahim, Kerstin Fischer, Makedonka Mitreva, Peter U. Fischer

## Abstract

Three molecularly undescribed filarial species were co-detected, while screening animals for *Brugia malayi*, the agent of lymphatic filariasis. Single microfilariae (Mf) isolated from blood samples of crab-eating macaques (*Macaca fascicularis*) from Belitung, Indonesia, and from pet dogs and cats in Sabah, Malaysia, were analyzed. Among 163 macaques, 33 (20.2%) were positive for large Mf (mean length 498.9 µm) similar to *Dirofilaria* (‘Belitung I’). One macaque was infected with small Mf (mean length 150.4 µm) (‘Belitung II’), with a high density of 17,150 Mf/mL. In two cats co-infected with *B. malayi*, Mf of a *Dirofilaria* species (‘Sabah’) with an average length of 299.1 µm were detected. Morphometric analysis of Mf showed distinct differences between these three species and other Mf described in the area. Whole genome amplification and genome sequencing of 24 individual Mf enabled phylogenetic analysis of mitochondrial genomes, and analysis of specific mitochondrial and nuclear barcode regions. The three Mf groups formed distinct clusters and did not match any currently available reference sequence. Cluster ‘Belitung I’ from macaques formed a sister group to all other *Dirofilaria*. Cluster ‘Belitung II’ included bird filariae and primate filariae of the genus *Mansonella* as close relatives. The cluster ‘Sabah’ formed a monophyletic group with the zoonotic species *D. asiatica* and *Dirofilaria* sp. ‘Thailand’. DNA of *Wolbachia* endobacteria was detected in Mf of ‘Belitung I’ and ‘Sabah’, but not in ‘Belitung II’. These findings highlight the limited understanding of filarial diversity in macaques and cats in Asia and underscore the need for a more comprehensive approach that combines morphological and molecular data to identify and assess the pathogenicity and zoonotic potential of these parasites.

**Author summary:** Filarial worms are parasitic nematodes that infect humans and animals and are often transmitted by the same vector mosquito. We identified three molecularly undescribed filarial species while investigating animals as reservoirs for the agent of lymphatic filariasis, *Brugia malayi* on Belitung Island, Indonesia, and in Sabah, Malaysia. Blood samples were collected from Indonesian macaques and Malaysian pet cats. Out of 163 macaques, 20.2% tested positive for exceptionally large microfilariae (Mf) of an unclassified *Dirofilaria*-like species (Belitung I’). Another filarial species (‘Belitung II’) with very small Mf, but with a remarkably high density of 17,150 Mf/mL was detected in one macaque. Two cats harbored medium sized Mf of a *Dirofilaria* species (Sabah’). Genetic analysis revealed unique phylogenetic clusters that did not match any reference sequence. *Dirofilaria* sp. ‘Sabah’ was closely related to the zoonotic *D. asiatica* complex, whereas ‘Belitung I’ clustered as a sister group to *Dirofilaria*. ‘Belitung II’ Mf clustered next to but not within the *Mansonella spp.* cluster. DNA of *Wolbachia* endobacteria was only detected in Mf of ‘Belitung I’ and ‘Sabah’. These findings highlight the limited understanding of filarial diversity in animals and underscore the need for a comprehensive approach that combines morphological and molecular data to identify and assess the pathogenicity and zoonotic potential of these parasites.

## Introduction

Filarial parasite species are globally distributed among mammalian, reptile, and avian hosts and are classified primarily under the superfamily Filarioidea, which is subdivided into Filariidae, Setariidae, Mesidionematidae, and Onchocercidae [1]. Despite their wide host range and diverse tropisms, many species remain poorly characterized. Morphological and molecular species description of members of this superfamily are often missing, and/or available morphological and molecular data cannot be reliably linked.

The limitations of relying solely on molecular data are illustrated by *Dirofilaria* sp. ‘hongkongensis’, which was first detected in a human in 2012 [2]. Although this species has been repeatedly reported in humans and animals in Bhutan, India [3–6], Sri Lanka [7], and Hong Kong [2, 8], its taxonomic placement remained unresolved for more than a decade. Coupling of the detailed morphological descriptions of adult worms and microfilariae (Mf) with genomic data enabled the species to be formally described as *Dirofilaria asiatica* sp. nov. [9]. *D. asiatica* sp. nov., *Dirofilaria* sp. ‘hongkongensis’, and *Dirofilaria* sp. ‘Thailand II’ form a closely related monophyletic group [10].

An accurate and complete species description of parasitic nematodes requires adult stages, particularly males, which are often accessible only through host necropsy. Although collecting and retrieving adult nematodes during surgery is feasible in domestic animals, obtaining comparable samples from wild animals, such as primates, is often impossible due to ethical and regulatory concerns. As a result, many filarial species circulating in wildlife remain insufficiently described, morphological and molecular data are not linked or both. This gap is concerning because several filarial species are of significant public health relevance. *Brugia malayi*, for example, causes lymphatic filariasis and is a target of the Global Program to Eliminate Lymphatic Filariasis of the World Health Organization. Other zoonotic species, such as *D. asiatica*, further underscore the importance of a One Health approach. Yet, despite their importance, filarial parasites have not been extensively studied within a One Health framework focused on human and animal health.

We recently demonstrated that non-human primates are an important reservoir for brugian filariasis. On Belitung Island, Indonesia, long-tailed macaques (*Macaca fascicularis*) were identified as the main reservoir hosts of *B. malayi* in an area endemic for human lymphatic filariasis [11]. These macaques are highly adaptable and synanthropic, in contrast to other primate species on the island that are largely nocturnal and avoid human contact [12]. Because macaques may harbor filarial species pathogenic to humans and mosquito vectors may bite both human and non-human primates, close interaction between hosts may facilitate transmission within species and between species [13, 14].

The objective of the present study was to provide molecular and morphometric data of Mf found in hosts of *B. malayi* in Southeast Asia to improve differential diagnosis. We examined non-human primates and domestic cats and dogs that live in close proximity with humans and are likely to be bitten by the same mosquito vectors. Whole mitochondrial genomes, together with mitochondrial and nuclear barcode regions, were analyzed to clarify phylogenetic relationships. These data expand existing molecular reference databases and enhance the ability to identify filarial species with zoonotic potential.

## Materials and methods

### Ethics approval and consent

The trapping and blood collection of animals were approved by the Ministries of Health and the Environment and Forestry of Indonesia (protocol #22-040365). The study received ethical approval from the ethical committee of Universitas Indonesia (no 515/UN2.F1/ETIK/PPM.00.02/2022) and was performed by veterinarians or, under their supervision, by veterinary technicians. Blood sample collection in Malaysia was approved by Animal Care and Use Committee (ACUC/KKM/02(01/2024)). Blood collection was performed by veterinarians.

### Study area and sample collection

Animal blood samples were collected as part of a study investigating the role of animal reservoirs for *B. malayi* in Indonesia and Malaysia. Belitung Island, Indonesia, is administratively divided into two districts, Belitung and Belitung Timur. Blood samples were obtained in five areas in the Belitung district (Selat Nasik and Petaling on Mendanau island, and Kembiri, Lassar, and Kacang Butor on the main island Belitung). The study area has been described in detail previously [15]. Macaques were trapped, and samples were collected at different time points (between 07:30-10:30am or between 3:00-7:30pm). The sample collection procedures have been described in detail previously [11]. In eastern Malaysia (Sabah, Borneo), pet cats and dogs were examined for Mf in an area with persistent *B. malayi* infection despite mass drug administration in the human population. Five blood samples that were morphologically positive for *B. malayi* were sent to Washington University in St. Louis for further molecular diagnostic confirmation.

Samples from two cats also contained Mf of another filarial species. Data on sex, village, sub-village, GPS coordinates, and habitat (forest near residence/ tourist attraction or palm oil plantation) were collected (S1 Table). None of the animals showed clinical signs of filarial infection.

### Morphological description

In Indonesia, two experienced microscopists examined the slides to identify and count Mf. A three-line thick blood smear (60 µl on the slide) was prepared following previously described procedures [16]. The slides were air-dried for 2 days at ambient temperature and then stained with 3% Giemsa solution (Sigma-Aldrich, Germany). Species identification was based on morphological features such as the presence of a sheath, the staining characteristics of the sheath, measurement of total body length and width, cephalic space (length, width and their ratio), the location of the nerve ring (NR), excretory pore (EP), excretory cell (EC), anal pore (AP), genital pore (GP), the “Innenkörper“, and the number and position of terminal nuclei (TN) in the tail structure of the microfilariae [17]. For morphometric comparison, measurements were taken from 15 individual Mf of the *Dirofilaria*-like unclassified Onchocercidae sp. ‘Belitung I’ from five macaques, 15 Mf of the *Mansonella*-like unclassified Onchocercidae sp. ‘Belitung II’ from a single macaque on Belitung Island, Indonesia, and 13 Mf of the unclassified *Dirofilaria* sp. ‘Sabah’ from one cat in Malaysia. Measurements included the length of the cephalic space clear of nuclei, the distance from the head to the nerve ring, the distance from the head to the excretory pore, the distance from the head to the anal pore, the distance from the head to the tail (terminal nuclei), the width at the level of the nerve ring, the total body length, and the ratio of mid-body width to the length of the nuclei-free cephalic space (Fig 1). All measurements and photographs were taken using an OLYMPUS BX40F4 microscope (Olympus Optical Co. Ltd., Japan) at 1000x magnification with cellSens Standard 1.18 software (Olympus Corporation of the Americas, USA).

**Fig 1.**
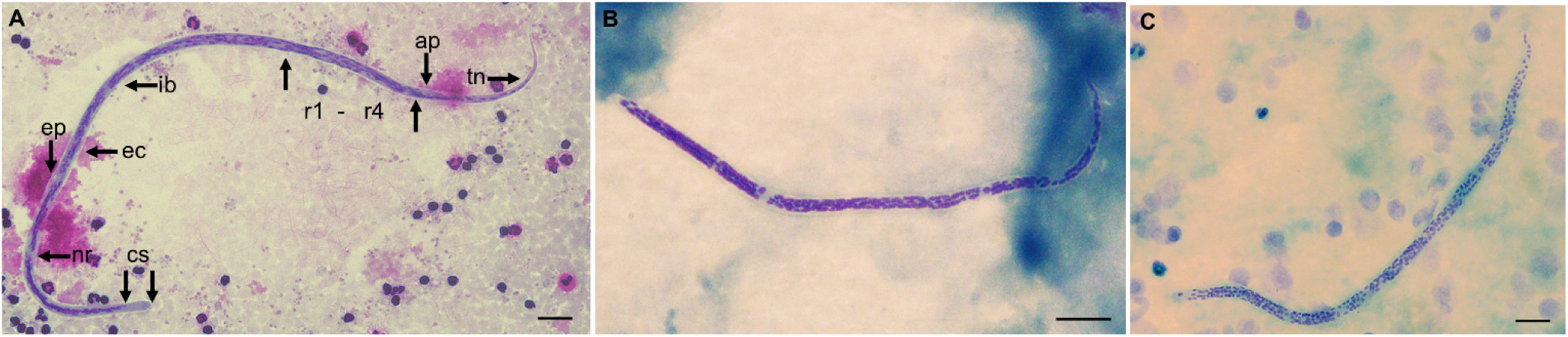
Morphological features of Giemsa-stained microfilaria. (A) Onchocercidae sp. ‘Belitung I’. (B) Onchocercidae sp. ‘Belitung II*’*. (C) *Dirofilaria* sp. ‘Sabah’. Abbreviation: cs-cephalic space; nr-nerve ring; ec-excretory cell; ep-excretory pore; ib-inner body or “Innenkörper”; ap-anal pore; r 1-4-rectal cells; tn-terminal nucleus, scale bar indicates 10 µm

### DNA extraction and sequencing

DNA was extracted from 50 µL of each blood sample, and the presence of pan-filarial DNA, specifically *B. malayi*, *B. pahangi,* and *D. immitis*, was assessed by qPCR as described in a previous study [11]. For selected samples that were morphologically microfilariae positive (S1 and S2 Tables), whole-genome amplification and sequencing were performed on individual Mf according to a previously described protocol [18]. Briefly, for DNA isolation from a single mf, we used a modified CGP DNA isolation protocol [18, 19]. To confirm a successful DNA isolation, a qPCR assay was performed to amplify a 28S ribosomal RNA pan-filarial fragment. qPCR conditions and primer pairs were described in previous studies [11, 20]. Positive samples were amplified using the Ready-To-Go GenomiPhi V3 DNA Amplification Kit (Cytiva, Marlborough, MA) according to the manufacturer’s recommendations. After whole-genome amplification, the sample was diluted 1:10, and the presence of parasite DNA was confirmed by qPCR. DNA concentration was measured by Qubit 4.0 (Thermo Fisher Scientific, Waltham, MA, USA) with the dsDNA BR Assay Kit (Thermo Fisher Scientific). A Kapa Hyper PCR-free library was generated from the amplified DNA and sequenced on Illumina’s NovaSeq platform (San Diego CA, USA, 2 × 150 bp paired-end reads) to ∼10 Gb per sample (S2 Table).

### Mitochondrial genome assembly and phylogenetic analyses

Sequencing reads were adapter- and quality-trimmed using Trimmomatic v0.39 [21]. GetOrganelle v1.7.5 [22] (-R 10 -k 21,45,65,85,105 -F animal_mt) was used to assemble *de novo* mitochondrial DNA from paired-end data. Mitochondrial genomes of *B. malayi* (AF538716) and *D. immitis* (AJ537512) were included in the seed database.

MITOS2 v2.1.9 [23] was used to determine the gene complement and the order of the protein-coding genes. The fully circularized assembled genomes and whole mitochondrial genome sequences of *Onchocercidae* spp. retrieved from GenBank were linearized to a common start position using rotate v1.0 [24]. Multiple sequence alignment was performed using MAFFT v7.505 [25] (--maxiterate 1000 --globalpair), followed by alignment trimming using trimAl v1.4 [26] (-automated1). A maximum likelihood phylogenetic tree was constructed using IQ-TREE2 v2.2.0 [27] (-B 10000 -bnni -alrt 10000 --robust-phy 0.98 -wsl) with *Spirocerca lupi* (KC305876) and *Tetrameres grusi* (MW648425) as outgroup taxa (-o).

### Cytochrome *c* oxidase I (COI) and 28S ribosomal RNA gene analyses

To increase the taxonomic resolution of phylogenetic analysis, the cytochrome *c* oxidase I (COI) region from the whole mitochondrial genome sequences of each Mf sample was used to search the NCBI database using BLASTn [28] and to retrieve reference sequences from the family *Onchocercidae* (NCBI:txid6296) (GenBank accessed on 5 November 2025) (S3 Table). *Tetrameres grusi* (MW648425) and *Spirocerca lupi* (KC305876) were included as outgroup. COI sequences were aligned using MAFFT v7.505 [25] and examined for correct codon alignment and in-frame translation using SeqKit translate [29]. Sequences shorter than 495 base pairs that resulted in incomplete coverage of the region were excluded. The final dataset included 141 sequences from 116 species belonging to 39 genera. A maximum likelihood phylogenetic tree was constructed using IQ-TREE2 v2.2.0 [27] (-B 10000 -bnni -alrt 10000 --robust-phy 0.98 -wsl -st CODON5) with a constrained tree search (-g). A whole mitochondrial genome tree was built as described above, excluding sequences from our unclassified Mf samples, and all nodes with bootstrap support below 99% were converted to multifurcating nodes using ggtree [30]. This tree was used as the constraint tree for the COI analysis.

The nuclear-encoded 28S rRNA gene was analyzed using a similar approach. Sequencing data from each Mf sample was assembled individually using SPAdes v4.2.0 [31]. The 28S rRNA region in each assembly was identified using minimap2 [32] with the 28S rRNA sequence from *Eufilaria acrocephalusi* (MT802308) as the query. The coordinates of the aligned region were converted to BED format and then to FASTA format using bedtools v2.31.0 [33]. Comparator sequences from *Onchocercidae* were retrieved from GenBank (S3 Table), aligned using MAFFT v7.505 [25] (--maxiterate 1000 --localpair), and trimmed using trimAl v1.4 [26] (-nogap). A maximum likelihood phylogenetic tree was generated using IQ-TREE2 v2.2.0 [27] as described above, except that *Setaria* spp. (EF199751, EF199750, KP760407, KP760408) were used as outgroup and no constraint tree was applied.

### Analysis of *Wolbachia* endosymbionts

Because *Wolbachia* copy numbers are lowest during the Mf stage [34], sequencing data were pooled per host animal (three to four Mf per host) to increase coverage prior to assembly using SPAdes v4.2.0 [31]. *Wolbachia*-derived sequences were identified in the resulting assemblies by performing dot plot analysis using D-Genies [35] with *Wolbachia* genomes from supergroups C, D, and F as target sequences. The *Wolbachia* surface protein (WSP) sequence was obtained by aligning WSP sequences from the *Wolbachia* endosymbiont of *O. volvulus* (CAL29428) and *D. immitis* (WP_175818468) to the assembly contigs using Exonerate [36] (--model protein2genome --bestn 1 --showtargetgff --showvulgar no --showalignment yes --refine region --refineboundary 10000). The corresponding protein sequence (translated CDS) was obtained from each hit using GffRead [37] (-y). Multiple sequence alignment was generated using MAFFT v7.505 [25] (--maxiterate 1000 --localpair) and trimmed using trimAl v1.4 [26] (-automated1). IQ-TREE2 v2.2.0 [27] was used for phylogenetic analysis as described above, with a *Wolbachia pipientis* reference WSP sequence (WP_250778599) as outgroup. Trees were visualized using the Interactive Tree of Life (iTOL) tool [38].

## Results

### Microscopy-based species identification, morphometric description, and density of microfilariae

In total, 33 of 163 (20.2%) blood samples from macaques contained Mf of a *Dirofilaria*-like unclassified Onchocercidae sp. (‘Belitung I’) (S1 Table). Among these, nine macaques (5.5%) were co-infected with *B. malayi* (S1 Table). The geometric mean Mf density for Onchocercidae sp. ‘Belitung I’ in macaques was 34 Mf/mL (range 17-484 Mf/mL) (S1 Table). In addition, *Mansonella*-like unclassified Onchocercidae sp. Mf (‘Belitung II’) collected from a male macaque in the forest habitat near a residence in the village of Selat Nasik could not be clearly identified by morphology. This macaque from Selat Nasik showed an exceptionally high Mf density of 17,150 Mf/mL. The arithmetic mean length of the Giemsa-stained Mf of ‘Belitung II’ was 150.4 ± 9.14 µm, whereas the *Dirofilaria*-like unclassified Onchocercidae sp. ‘Belitung I’ was more than three times larger, with a head-to-tail length of 498.9 ± 13.15 µm. Out of the five samples obtained from Malaysia (three cats and two dogs), two cats harbored *Dirofilaria* sp. ‘Sabah’ Mf. The head-to-tail length was 299.1 ± 6.52 µm (Table 1). The Mf density of this species was not determined. No sheaths and terminal nuclei were detected in any of the analyzed Mf.

**Table 1.**
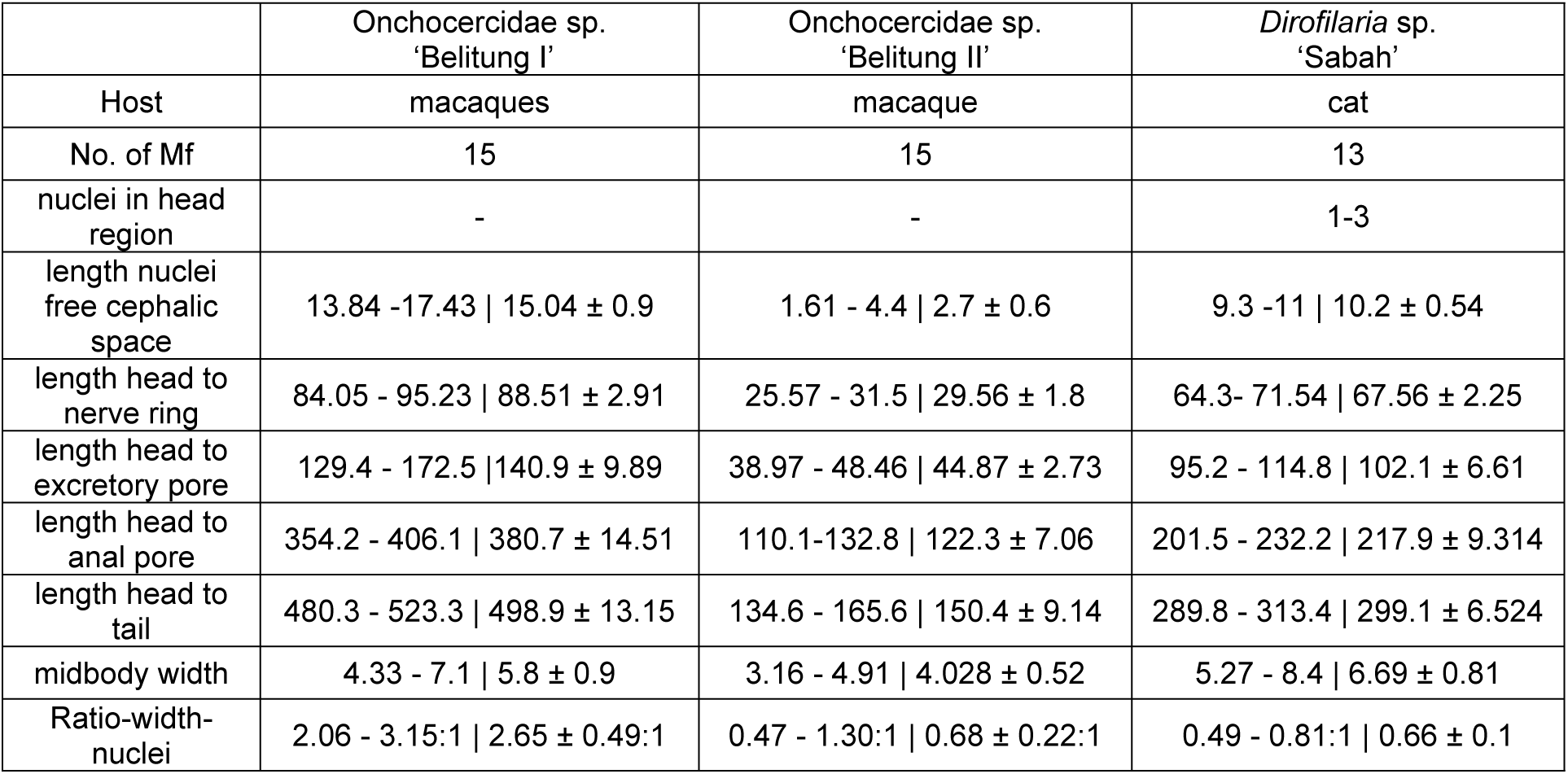
Morphometry of Giemsa-stained microfilariae. Measurements (in µm) are presented as range and arithmetic mean ± SD.

|  | Onchocercidae sp.<br>'Belitung I' | Onchocercidae sp.<br>'Belitung II' | *Dirofilaria* sp.<br>'Sabah' |
| --- | --- | --- | --- |
| Host | macaques | macaque | cat |
| No. of Mf | 15 | 15 | 13 |
| nuclei in head region | - | - | 1-3 |
| length nuclei free cephalic space | 13.84 - 17.43 \| 15.04 ± 0.9 | 1.61 - 4.4 \| 2.7 ± 0.6 | 9.3 - 11 \| 10.2 ± 0.54 |
| length head to nerve ring | 84.05 - 95.23 \| 88.51 ± 2.91 | 25.57 - 31.5 \| 29.56 ± 1.8 | 64.3 - 71.54 \| 67.56 ± 2.25 |
| length head to excretory pore | 129.4 - 172.5 \| 140.9 ± 9.89 | 38.97 - 48.46 \| 44.87 ± 2.73 | 95.2 - 114.8 \| 102.1 ± 6.61 |
| length head to anal pore | 354.2 - 406.1 \| 380.7 ± 14.51 | 110.1 - 132.8 \| 122.3 ± 7.06 | 201.5 - 232.2 \| 217.9 ± 9.314 |
| length head to tail | 480.3 - 523.3 \| 498.9 ± 13.15 | 134.6 - 165.6 \| 150.4 ± 9.14 | 289.8 - 313.4 \| 299.1 ± 6.524 |
| midbody width | 4.33 - 7.1 \| 5.8 ± 0.9 | 3.16 - 4.91 \| 4.028 ± 0.52 | 5.27 - 8.4 \| 6.69 ± 0.81 |
| Ratio-width-nuclei | 2.06 - 3.15:1 \| 2.65 ± 0.49:1 | 0.47 - 1.30:1 \| 0.68 ± 0.22:1 | 0.49 - 0.81:1 \| 0.66 ± 0.1 |

### Diagnostic qPCR results

The qPCR results for filarial species originating from macaques have been previously published [11]. None of the macaques tested positive for *B. pahangi* or *D. immitis*. For the Malaysian sample, all samples were pan-filarial positive. Three cat samples were positive for *B. malayi* and *B. pahangi*, including both cats that contained *Dirofilaria* sp. ‘Sabah’. None of the cat samples were positive for *D. immitis*. Only one dog tested positive for *D. immitis* (S1 Table).

### Phylogenetic analyses based on whole mitochondrial genome

Mitochondrial genomes were *de novo* assembled for each Mf sample, yielding complete circularized genomes within the expected size range (13.6-13.8 kb) and displaying the conserved set of 12 protein-coding genes and the characteristic gene order of filarial nematodes [39]. To determine the taxonomic placement of the unclassified filarial species, these genomes were compared with mitochondrial sequences of other Onchocercidae taxa available in GenBank (S3 Table). Pairwise nucleotide identity analyses showed that Onchocercidae sp. ‘Belitung I’ was most similar to *Dirofilaria repens* (KX265048; 88.0% identity; 98% query coverage). Onchocercidae sp. ‘Belitung II’ showed the highest similarity also to *D. repens* (KX265048; 83.3% identity; 98% query coverage), whereas *Dirofilaria* sp. ‘Sabah’ most closely matched *D. asiatica* (PQ131197; 96.9% identity; 100% query coverage).

A maximum likelihood phylogenetic tree including 23 additional Onchocercidae taxa revealed that Onchocercidae sp. ‘Belitung I’ formed a well-supported clade outside the *Dirofilaria* genus and appeared as its sister group, suggesting that it may represent a distinct lineage (Fig 2). In contrast, *Dirofilaria* sp. ‘Sabah’ clustered within the *Dirofilaria* genus and grouped monophyletically with *D. asiatica* and *Dirofilaria* sp. ‘Thailand II’ [9, 10]. The *Mansonella*-like Onchocercidae sp. ‘Belitung II’ formed a separate clade that was the sister group to the genus *Mansonella*.

**Fig 2.**
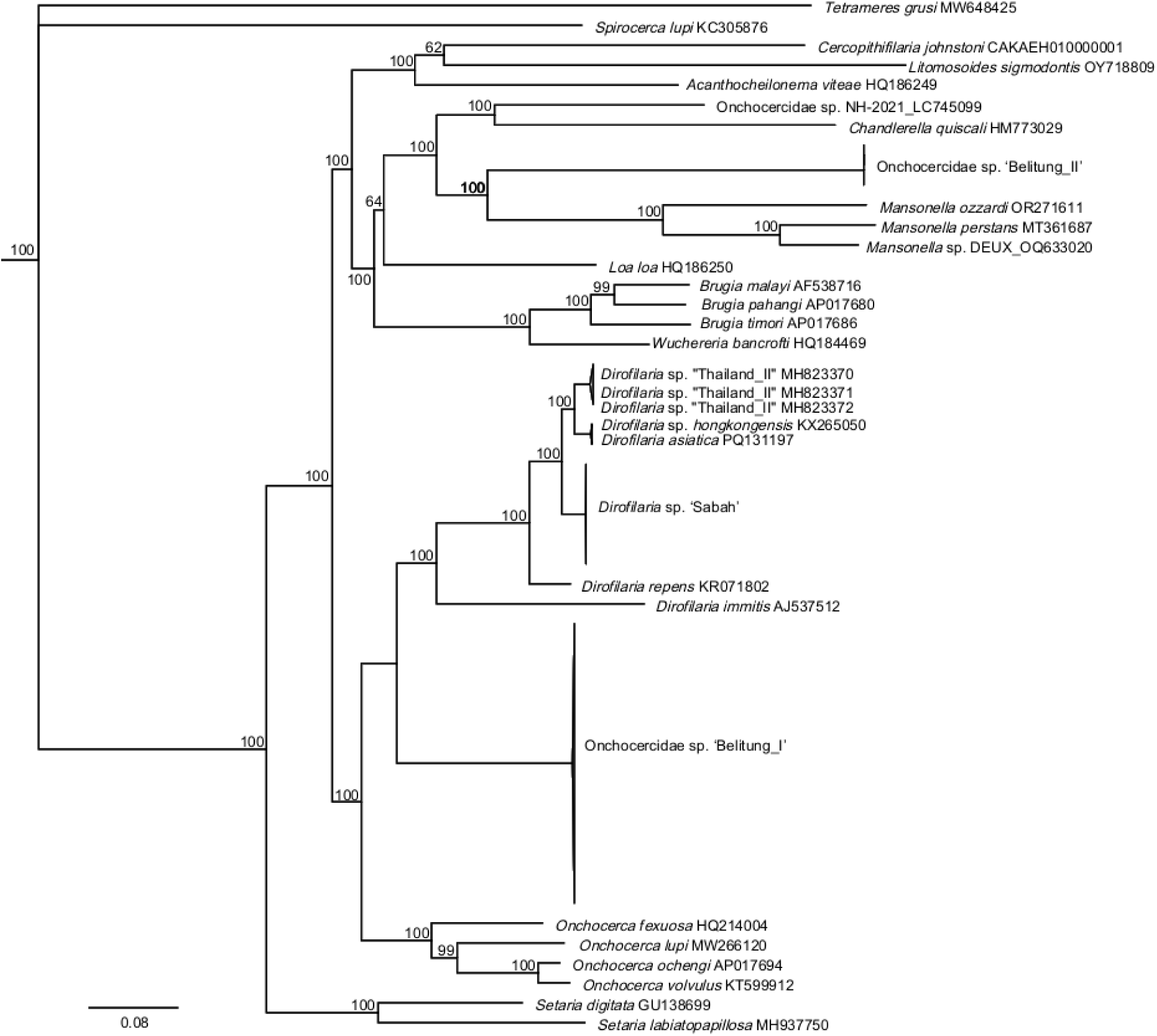
Whole mitochondrial genome maximum likelihood phylogenetic tree. The scale bar represents 0.08 substitutions per site and node support was obtained by ultrafast bootstrap approximation [40].

### Phylogenetic analyses based on cytochrome *c* oxidase I (COI) and 28S ribosomal RNA gene

To improve taxonomic resolution, the COI region from each mitochondrial genome was queried against the NCBI database to retrieve reference sequences from the family Onchocercidae (NCBI:txid6296). The COI sequences of Onchocercidae sp. ‘Belitung I’ showed the highest similarity to an unclassified *D. repens*-like species from Georgia, USA (PQ191455; 94.9% identity; 100% query coverage). The closest match to Onchocercidae sp. ‘Belitung II’ COI sequences were *Eufilaria sylviae* (MT800771; 89.7% identity; 100% query coverage), identified from a garden warbler (*Sylvia borin*) in Lithuania. The COI sequences of *Dirofilaria* sp. ‘Sabah’ were most similar to *D. asiatica* (PV523835; 97.4% identity; 100% query coverage), obtained from a dog in Sri Lanka.

Sequence similarity searches using the nuclear-encoded 28S rRNA gene indicated that Onchocercidae sp. ‘Belitung I’ was most similar to *Dirofilaria ursi* (PV389592; 95.9% identity; 100% query coverage), obtained from a Japanese black bear in Japan. The 28S rRNA gene sequences of Onchocercidae sp. ‘Belitung II’ most closely matched *Mansonella perstans* (MN432520; 90.3% identity; 100% query coverage), obtained from a human in Brazil. *Dirofilaria* sp. ‘Sabah’ showed the highest similarity to *D. repens* (KP760376; 98.7% identity; 100% query coverage), obtained from a dog in Italy.

Maximum-likelihood phylogenies based on COI and 28S rRNA sequences placed Onchocercidae spp. Belitung I’ and ‘Belitung II’ in positions consistent with those inferred from the whole mitochondrial genome analyses (Figs 3 and 4). The COI sequences of *Dirofilaria* sp. ‘Sabah’ also fell within the clade formed by the genus *Dirofilaria*. In this analysis, however, they were monophyletic with *Dirofilaria* sp. ‘Thailand II’ and paraphyletic to *D. asiatica*, which differed from the topology observed in the whole mitochondrial genome tree (Fig 2). Because 28S rRNA sequences for *D. asiatica* and *Dirofilaria* sp. ‘Thailand II’ were not available, their relationships could not be evaluated using nuclear genetic markers.

**Fig 3.**
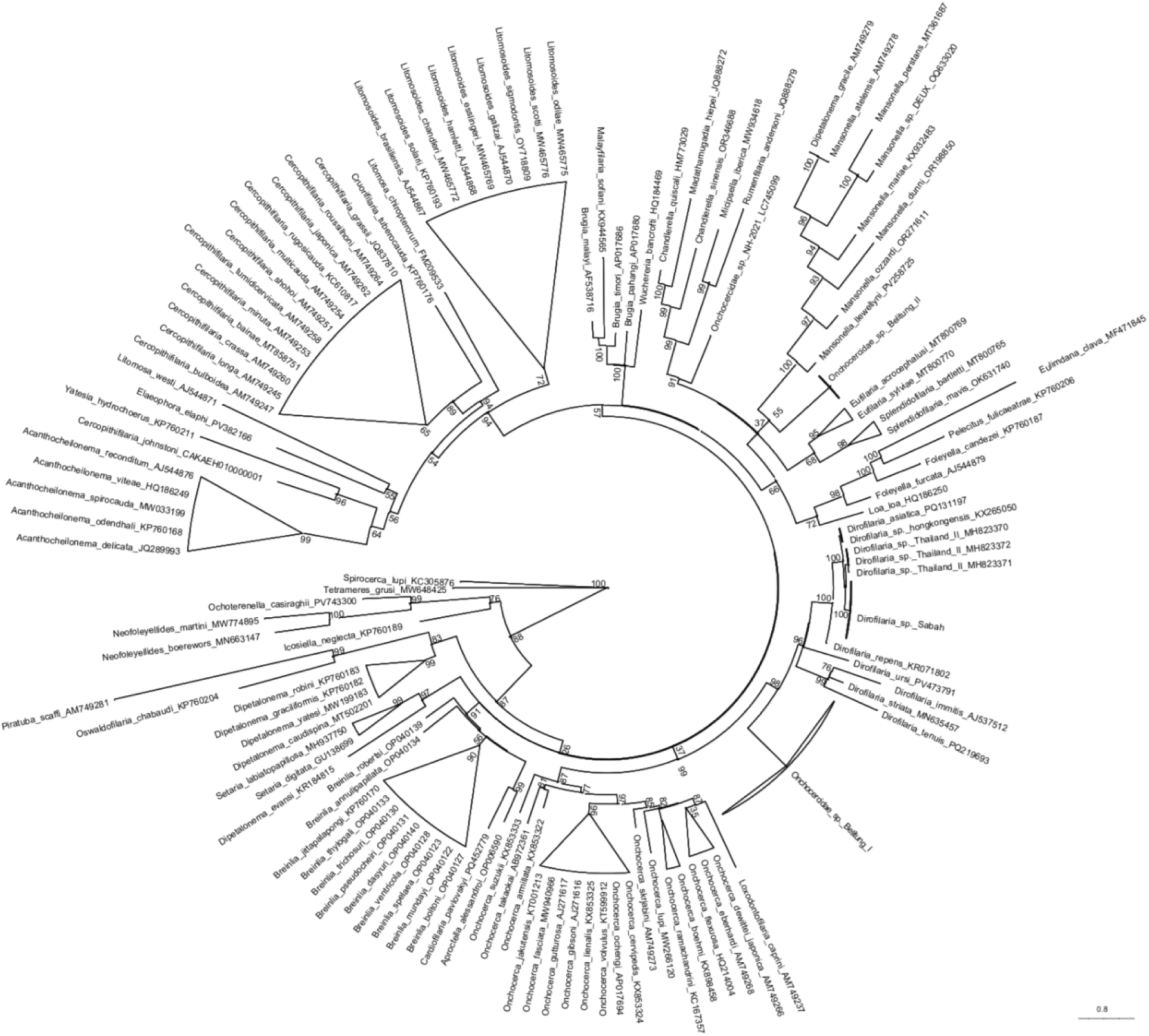
Cytochrome *c* oxidase I maximum likelihood phylogenetic tree. The scale bar represents 0.8 substitutions per site and node support was obtained by ultrafast bootstrap approximation [40].

**Fig 4.**
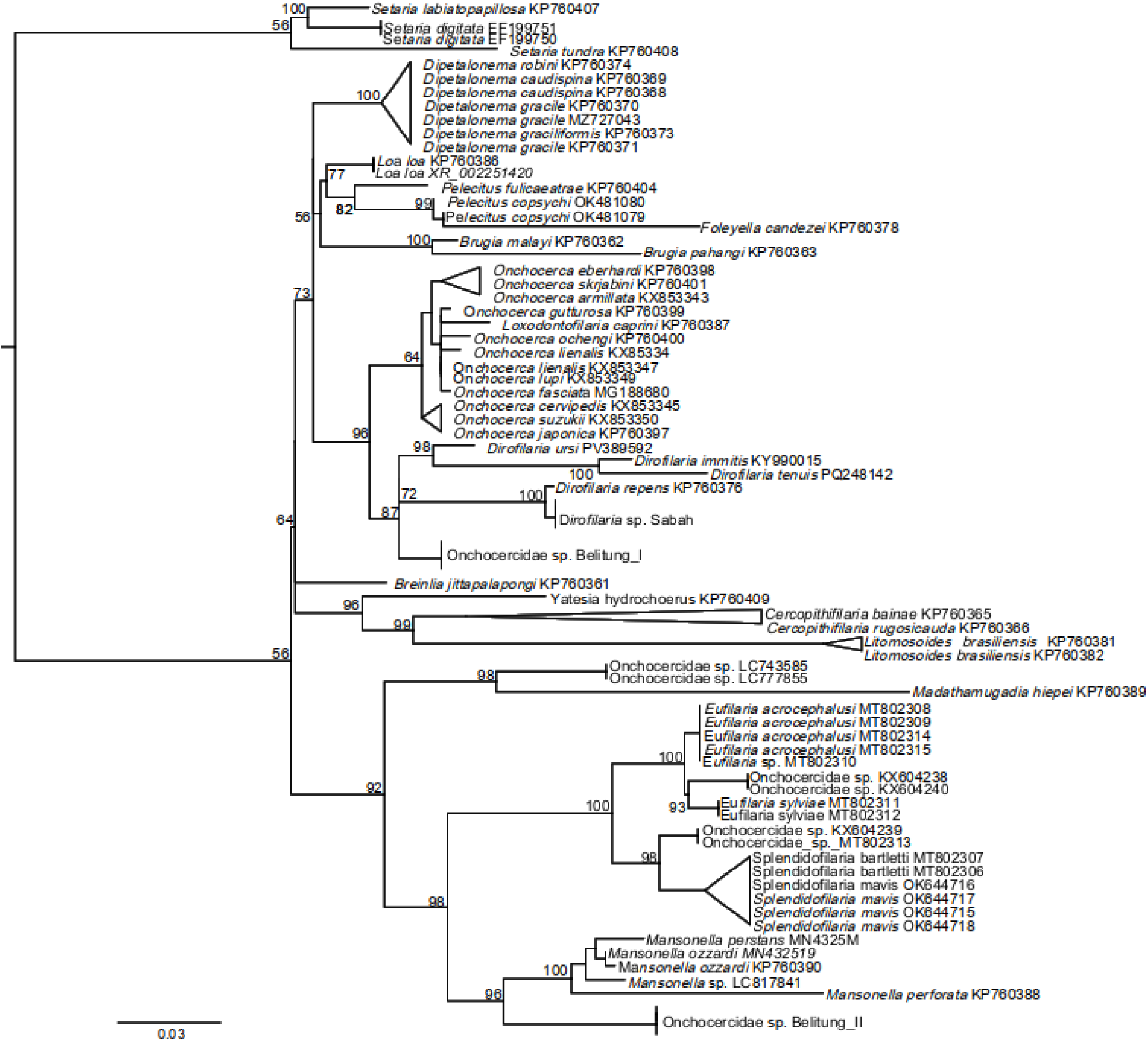
28S ribosomal RNA maximum likelihood phylogenetic tree. The scale bar represents 0.03 substitutions per site and node support was obtained by ultrafast bootstrap approximation [40].

### Presence and absence of *Wolbachia* endosymbiont

Contigs assembled from Onchocercidae sp. ‘Belitung I’, Onchocercidae sp. ‘Belitung II’ and *Dirofilaria* sp. ‘Sabah’ were screened for *Wolbachia* DNA sequences using dot plot analyses against representative genomes from supergroups C, D, and F. *Wolbachia* DNA was detected in Onchocercidae sp. ‘Belitung I’ and *Dirofilaria* sp. ‘Sabah’, but not in Onchocercidae sp. ‘Belitung II’ (S1 Fig).

Phylogenetic analysis of the *Wolbachia* surface protein (WSP) placed both detected endosymbionts in supergroup C (Fig 5). The *Wolbachia* from Onchocercidae sp. ‘Belitung I’ most closely matched the endosymbiont of *Onchocerca gibsoni* (WP_275601952; 89.9 % identity; 98 % query coverage), while the *Wolbachia* from Dirofilaria sp. ‘Sabah’ was most similar to that of *Dirofilaria asiatica* (WLD16215; 99.1 % identity; 94 % query coverage).

**Fig 5.**
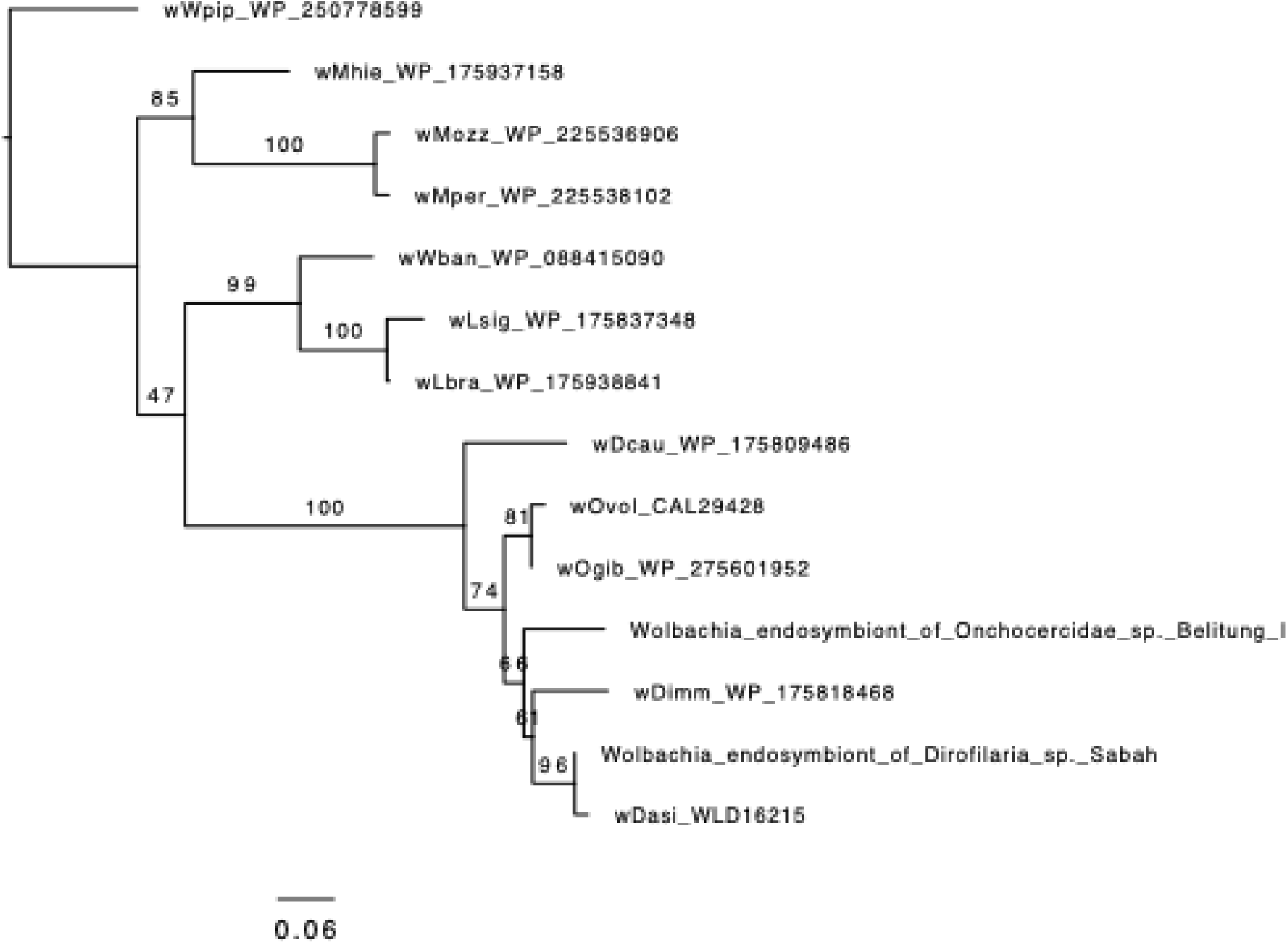
*Wolbachia* surface protein (WSP) maximum likelihood phylogenetic tree. Node support was obtained by ultrafast bootstrap approximation [40].

## Discussion

This study provides important morphometric and molecular characterizations of hitherto undescribed Mf found in reservoir hosts of *B. malayi* in Indonesia and Malaysia to facilitate differential diagnostics. Complete mitochondrial genomes, nuclear barcode regions, and *Wolbachia* marker genes were assembled and used in phylogenetic analyses that clarified the taxonomic placement of these previously uncharacterized filarial species within the family Onchocercidae and the genus *Dirofilaria*. These new molecular data highlight the extent of undescribed genetic and species diversity within Onchocercidae and expand public sequence databases that support molecular diagnostics.

The Onchocercidae sp. ‘Belitung I’ Mf were detected by microscopy in approximately 20% of the macaques. With an average length of around 500 µm, they are longer than any Mf known from humans and most commonly reported Mf from animals. To our knowledge, no molecularly described Mf match this size range. However, within the genus *Dirofilaria*, one species, *D. magnilarvatum*, was named to reflect the unusually large size of its unsheathed Mf (580.9 ± 10 µm) [41, 42]. This species, which was described from *Macaca irus* (now *Macaca fascicularis*) in Malaysia, shows no periodicity and tends to accumulate in arterioles in the skin and tail. Because our Mf were found in the same host species, and natural or technical size variation is common, it is possible that ‘Belitung I’ corresponds to *D. magnilarvatum*. However, the absence of molecular data for *D. magilavatum* prevents definitive identification. Reported vectors for this species include *Mansonia bonneae, M. annulata* [43], *M. longipalpis,* and *M. uniformis* [44], which are also known vectors of *B. malayi*, that was also detected in these macaques [11]. A previous study in the same area reported Mf of a *Dirofilaria sp.* in macaques that may correspond to the species identified in our samples, although no morphological and molecular description were provided [45]. Taken together, these findings suggest that large *Dirofilaria-*like Mf are relatively common in macaques on Belitung Island. Because parasites circulating in non-human primates may have zoonotic potential, further investigation of this lineage is warranted.

*Dirofilaria* sp. ‘Sabah’ morphologically resembled *D. asiatica* Mf. The Mf of both species had very similar lengths (299.1 µm vs. 300.3 µm), and the number of nuclei in the head space was also comparable (1-3 vs. 2-3) [9]. However, because different staining methods were used, direct comparison should be interpreted with caution. The close relationship of the ‘Belitung I’ and particularly the ‘Sabah’ Mf to *D. asiatica* and *Dirofilaria* sp. ‘Thailand II’ suggests substantial diversity within the genus *Dirofilaria* in Asia [9, 10, 46]. Weather *Dirofilaria* sp. ‘Sabah’ represents a cryptic species, or a novel species can only be determined through the collection of adult worms and additional molecular data. Given its close phylogenetic proximity to *D. asiatica*, it is plausible that *Dirofilaria* sp. ‘Sabah’ shares similar epidemiological and ecological characteristics, including a zoonotic potential.

The Mf of Onchocercidae sp. ‘Belitung II’ are similar in size to those of other *Mansonella* species [47]. Although whole mitochondrial genomes provide more robust phylogenetic information, their utility is limited because complete mitochondrial sequences are unavailable for many Filarioidea species. In our analyses, ‘Belitung II’ clustered as a sister group to *Mansonella*, a genus commonly found in humans and primates in Africa and South America [48, 49]. The only *Mansonella* species described from Asia is *M. dunni*, reported from the common tree shrew (*Tupaia glis*) [50]. Because molecular data for non-human primate *Mansonella* species are lacking, other filarial species previously described in macaques, such as *Edesonfilaria malayensis* or *Macacanema formosana*, cannot be excluded. For example, the Mf of *M. formosana* found in Taiwanese macaques (*Macaca cyclopis*) measured 152 µm [51], closely resembling the length of ‘Belitung II’ (150 µm). Of particular interest is the low prevalence but extremely high Mf density (17,150 Mf/mL) of ‘Belitung II’, which may indicate highly efficient transmission by a specific vector species or immunosuppressive conditions in the affected macaque.

The *Wolbachia* profiles of the newly identified filarial species further support their inferred taxonomic relationships. *Wolbachia* from Onchocercidae sp. ‘Belitung I’ and *Dirofilaria* sp. ‘Sabah’ clustered within supergroup C and closely matched endosymbionts of *Onchocerca* and *Dirofilaria* species, findings that are broadly consistent with mitochondrial and nuclear phylogenies. In contrast, the absence of detectable *Wolbachia* in Onchocercidae sp. ‘Belitung II’ aligns with its placement near the *Wolbachia*-free species *Chandlerella quiscali* [52], although low infection levels may account for the negative results, as has been reported for other *Mansonella* species [53].

For all three molecularly undescribed species in this study, comprehensive information on critical aspects such as pathogenicity, vector competence, host preference, and geographical distribution is lacking. This absence of data represents a major gap in our understanding of filarial biocoenoses and underscores the need for further investigation.

## Conclusion

Analysis of individual Mf provides an effective approach for linking morphological features with genome data. Our findings highlight the limited knowledge of filarial species diversity in non-human primates and companion animals, some of which may have zoonotic potential and/or aggravate diagnosis of *B. malayi* infection.

## Supporting information

Table S1: Metadata, microscopy, and qPCR results

Table S2: NCBI accession numbers for sequences generated in this study Table S3: NCBI accession numbers for comparator sequences

Figure S1: Dot plots comparing assembly scaffolds (query) with *Wolbachia* genomes of Onchocercidae spp. (target).

## Acknowledgments

We thank Mr. Sudirman and the rest of the field team of the Universitas Indonesia, and the Local Health Authority, the Primary Health Centers and Health Polytechnic of the Ministry of Health in Belitung District for their expert technical support. In addition, we would like to thank all pet owners for their support.

## References

1. Orihel TC, Eberhard ML. Zoonotic Filariasis. Clinical Microbiology Reviews. 1998;11(2):366-81. doi: doi:10.1128/cmr.11.2.366.

2. To KK, Wong SS, Poon RW, Trendell-Smith NJ, Ngan AH, Lam JW, et al. A novel Dirofilaria species causing human and canine infections in Hong Kong. J Clin Microbiol. 2012;50(11):3534–41. Epub 20120822. doi: 10.1128/jcm.01590-12. PubMed PMID: 22915604; PubMed Central PMCID: PMCPMC3486214.

3. Gowrishankar S, Aravind M, Sastya S, Latha BR, Azhahianambi P, Vairamuthu S, et al. Dirofilaria hongkongensis - A first report of potential zoonotic dirofilariosis infection in dogs from Tamil Nadu. Vet Parasitol Reg Stud Reports. 2019;18:100326. Epub 20190806. doi: 10.1016/j.vprsr.2019.100326. PubMed PMID: 31796197.

4. Kumar A, Sreedhar A, Biswas L, Prabhat S, Suresh P, Asokan A, et al. Candidatus Dirofilaria Hongkongensis Infections in Humans During 2005 to 2020, in Kerala, India. Am J Trop Med Hyg. 2021;104(6):2046–9. Epub 20210412. doi: 10.4269/ajtmh.20-1521. PubMed PMID: 33844649; PubMed Central PMCID: PMCPMC8176466.

5. Winkler S, Pollreisz A, Georgopoulos M, Bagò-Horvath Z, Auer H, To KK, et al. Candidatus Dirofilaria hongkongensis as Causative Agent of Human Ocular Filariosis after Travel to India. Emerg Infect Dis. 2017;23(8):1428–31. doi: 10.3201/eid2308.170423. PubMed PMID: 28726623; PubMed Central PMCID: PMCPMC5547781.

6. Schroeder J, Rothe C, Hoerauf A, Kroidl I, Pfarr K, Hübner MP. First case of Dirofilaria hongkongensis infection in Germany presenting as a breast tumour. J Travel Med. 2023;30(8). doi: 10.1093/jtm/taad121. PubMed PMID: 37738591; PubMed Central PMCID: PMCPMC10755165.

7. Atapattu U, Koehler AV, Huggins LG, Wiethoelter A, Traub RJ, Colella V. Dogs are reservoir hosts of the zoonotic Dirofilaria sp. ’hongkongensis’ and potentially of Brugia sp. Sri Lanka genotype in Sri Lanka. One Health. 2023;17:100625. Epub 20230830. doi: 10.1016/j.onehlt.2023.100625. PubMed PMID: 38024272; PubMed Central PMCID: PMCPMC10665175.

8. Manathunga T, Tse M, Perles L, Beugnet F, Barrs V, Otranto D. Zoonotic Dirofilaria sp. "hongkongensis" in subcutaneous nodules from dogs and cats, Hong Kong SAR. Parasit Vectors. 2024;17(1):469. Epub 20241115. doi: 10.1186/s13071-024-06544-7. PubMed PMID: 39548498; PubMed Central PMCID: PMCPMC11566833.

9. Colella V, Young ND, Manzanell R, Atapattu U, Sumanam SB, Huggins LG, et al. Dirofilaria asiatica sp. nov. (Spirurida: Onchocercidae) – Defined using a combined morphological-molecular approach. International Journal for Parasitology. 2025;55(8):461–74. doi: 10.1016/j.ijpara.2025.04.006.

10. Yilmaz E, Fritzenwanker M, Pantchev N, Lendner M, Wongkamchai S, Otranto D, et al. The Mitochondrial Genomes of the Zoonotic Canine Filarial Parasites Dirofilaria (Nochtiella) repens and Candidatus Dirofilaria (Nochtiella) hongkongensis Provide Evidence for Presence of Cryptic Species. PLoS Negl Trop Dis. 2016;10(10):e0005028. Epub 20161011. doi: 10.1371/journal.pntd.0005028. PubMed PMID: 27727270; PubMed Central PMCID: PMCPMC5058507.

11. Diekmann I, Supali T, Fischer K, Iskandar E, Sugianto N, Destani Y, et al. Brugia malayi and other filarial parasite species in animals in areas endemic for lymphatic filariasis in Belitung District, Indonesia [version 1]. VeriXiv. 2025;2(55). doi: 10.12688/verixiv.849.1.

12. Hasan MU, Arismayanti E, Perwitasari-Farajallah D, Tsuji Y, Widayati Kanthi A. Behavioural responses of long-tailed macaques (Macaca fascicularis) to environmental fluctuations: a preliminary study on Belitung Island, Indonesia. Folia Primatologica. 2025:1–20. doi: 10.1163/14219980-bja10064.

13. Gardner MB, Luciw PA. Macaque models of human infectious disease. Ilar j. 2008;49(2):220–55. doi: 10.1093/ilar.49.2.220. PubMed PMID: 18323583; PubMed Central PMCID: PMCPMC7108592.

14. Pedersen AB, Davies TJ. Cross-species pathogen transmission and disease emergence in primates. Ecohealth. 2009;6(4):496–508. Epub 20100316. doi: 10.1007/s10393-010-0284-3. PubMed PMID: 20232229; PubMed Central PMCID: PMCPMC7087625.

15. Supali T, Djuardi Y, Santoso, Sianipar LR, Suryaningtyas NH, Alfian R, et al. Surveillance and Selective Treatment of Brugia malayi Filariasis Eleven Years after Stopping Mass Drug Administration in Belitung District, Indonesia. Am J Trop Med Hyg. 2024;110(1):111–6. Epub 20231127. doi: 10.4269/ajtmh.23-0255. PubMed PMID: 38011734; PubMed Central PMCID: PMCPMC10793014.

16. Supali T, Djuardi Y, Lomiga A, Nur Linda S, Iskandar E, Goss CW, et al. Comparison of the Impact of Annual and Semiannual Mass Drug Administration on Lymphatic Filariasis Prevalence in Flores Island, Indonesia. Am J Trop Med Hyg. 2019;100(2):336–43. doi: 10.4269/ajtmh.18-0570. PubMed PMID: 30560772; PubMed Central PMCID: PMCPMC6367633.

17. Tongu Y. Ultrastructural studies on the microfilaria of Brugia malayi. Acta medica Okayama. 1974;28 3:219–42.

18. Choi YJ, Fischer K, Méité A, Koudou BG, Fischer PU, Mitreva M. Distinguishing recrudescence from reinfection in lymphatic filariasis. EBioMedicine. 2024;105:105188. Epub 20240607. doi: 10.1016/j.ebiom.2024.105188. PubMed PMID: 38848649; PubMed Central PMCID: PMCPMC11200287.

19. Moore L, Leongamornlert D, Coorens THH, Sanders MA, Ellis P, Dentro SC, et al. The mutational landscape of normal human endometrial epithelium. Nature. 2020;580(7805):640–6. Epub 20200422. doi: 10.1038/s41586-020-2214-z. PubMed PMID: 32350471.

20. Laidoudi Y, Marie JL, Tahir D, Watier-Grillot S, Mediannikov O, Davoust B. Detection of Canine Vector-Borne Filariasis and Their Wolbachia Endosymbionts in French Guiana. Microorganisms. 2020;8(5). Epub 20200521. doi: 10.3390/microorganisms8050770. PubMed PMID: 32455576; PubMed Central PMCID: PMCPMC7285362.

21. Bolger AM, Lohse M, Usadel B. Trimmomatic: a flexible trimmer for Illumina sequence data. Bioinformatics. 2014;30(15):2114–20. Epub 2014/04/04. doi: 10.1093/bioinformatics/btu170. PubMed PMID: 24695404; PubMed Central PMCID: PMCPMC4103590.

22. Jin JJ, Yu WB, Yang JB, Song Y, dePamphilis CW, Yi TS, et al. GetOrganelle: a fast and versatile toolkit for accurate de novo assembly of organelle genomes. Genome Biol. 2020;21(1):241. Epub 2020/09/12. doi: 10.1186/s13059-020-02154-5. PubMed PMID: 32912315; PubMed Central PMCID: PMCPMC7488116.

23. Donath A, Juhling F, Al-Arab M, Bernhart SH, Reinhardt F, Stadler PF, et al. Improved annotation of protein-coding genes boundaries in metazoan mitochondrial genomes. Nucleic Acids Res. 2019;47(20):10543–52. Epub 2019/10/05. doi: 10.1093/nar/gkz833. PubMed PMID: 31584075; PubMed Central PMCID: PMCPMC6847864.

24. Durbin R, De Sanctis B, Blumer M. Rotate: A command-line program to rotate circular DNA sequences to start at a given position or string. Wellcome Open Res. 2023;8:401. Epub 2024/04/29. doi: 10.12688/wellcomeopenres.19568.1. PubMed PMID: 38680652; PubMed Central PMCID: PMCPMC11056001.

25. Katoh K, Standley DM. MAFFT multiple sequence alignment software version 7: improvements in performance and usability. Mol Biol Evol. 2013;30(4):772–80. Epub 20130116. doi: 10.1093/molbev/mst010. PubMed PMID: 23329690; PubMed Central PMCID: PMCPMC3603318.

26. Capella-Gutierrez S, Silla-Martinez JM, Gabaldon T. trimAl: a tool for automated alignment trimming in large-scale phylogenetic analyses. Bioinformatics. 2009;25(15):1972–3. Epub 2009/06/10. doi: 10.1093/bioinformatics/btp348. PubMed PMID: 19505945; PubMed Central PMCID: PMCPMC2712344.

27. Nguyen L-T, Schmidt HA, von Haeseler A, Minh BQ. IQ-TREE: A Fast and Effective Stochastic Algorithm for Estimating Maximum-Likelihood Phylogenies. Molecular Biology and Evolution. 2014;32(1):268–74. doi: 10.1093/molbev/msu300.

28. Altschul SF, Gish W, Miller W, Myers EW, Lipman DJ. Basic local alignment search tool. J Mol Biol. 1990;215(3):403–10. doi: 10.1016/s0022-2836(05)80360-2. PubMed PMID: 2231712.

29. Shen W, Le S, Li Y, Hu F. SeqKit: A Cross-Platform and Ultrafast Toolkit for FASTA/Q File Manipulation. PLoS One. 2016;11(10):e0163962. Epub 2016/10/06. doi: 10.1371/journal.pone.0163962. PubMed PMID: 27706213; PubMed Central PMCID: PMCPMC5051824.

30. Yu G, Lam TT, Zhu H, Guan Y. Two Methods for Mapping and Visualizing Associated Data on Phylogeny Using Ggtree. Mol Biol Evol. 2018;35(12):3041–3. Epub 2018/10/24. doi: 10.1093/molbev/msy194. PubMed PMID: 30351396; PubMed Central PMCID: PMCPMC6278858.

31. Bankevich A, Nurk S, Antipov D, Gurevich AA, Dvorkin M, Kulikov AS, et al. SPAdes: a new genome assembly algorithm and its applications to single-cell sequencing. J Comput Biol. 2012;19(5):455–77. Epub 2012/04/18. doi: 10.1089/cmb.2012.0021. PubMed PMID: 22506599; PubMed Central PMCID: PMCPMC3342519.

32. Li H. Minimap2: pairwise alignment for nucleotide sequences. Bioinformatics. 2018;34(18):3094–100. Epub 2018/05/12. doi: 10.1093/bioinformatics/bty191. PubMed PMID: 29750242; PubMed Central PMCID: PMCPMC6137996.

33. Quinlan AR, Hall IM. BEDTools: a flexible suite of utilities for comparing genomic features. Bioinformatics. 2010;26(6):841–2. Epub 2010/01/30. doi: 10.1093/bioinformatics/btq033. PubMed PMID: 20110278; PubMed Central PMCID: PMCPMC2832824.

34. McGarry HF, Egerton GL, Taylor MJ. Population dynamics of Wolbachia bacterial endosymbionts in Brugia malayi. Mol Biochem Parasitol. 2004;135(1):57–67. Epub 2004/08/04. doi: 10.1016/j.molbiopara.2004.01.006. PubMed PMID: 15287587.

35. Cabanettes F, Klopp C. D-GENIES: dot plot large genomes in an interactive, efficient and simple way. PeerJ. 2018;6:e4958. Epub 2018/06/12. doi: 10.7717/peerj.4958. PubMed PMID: 29888139; PubMed Central PMCID: PMCPMC5991294.

36. Slater GS, Birney E. Automated generation of heuristics for biological sequence comparison. BMC Bioinformatics. 2005;6:31. Epub 2005/02/17. doi: 10.1186/1471-2105-6-31. PubMed PMID: 15713233; PubMed Central PMCID: PMCPMC553969.

37. Pertea G, Pertea M. GFF Utilities: GffRead and GffCompare. F1000Res. 2020;9. Epub 2020/09/22. doi: 10.12688/f1000research.23297.2. PubMed PMID: 32489650; PubMed Central PMCID: PMCPMC7222033.

38. Letunic I, Bork P. Interactive Tree of Life (iTOL) v6: recent updates to the phylogenetic tree display and annotation tool. Nucleic Acids Research. 2024;52(W1):W78–W82. doi: 10.1093/nar/gkae268.

39. McNulty SN, Mullin AS, Vaughan JA, Tkach VV, Weil GJ, Fischer PU. Comparing the mitochondrial genomes of Wolbachia-dependent and independent filarial nematode species. BMC Genomics. 2012;13:145. Epub 2012/04/26. doi: 10.1186/1471-2164-13-145. PubMed PMID: 22530989; PubMed Central PMCID: PMCPMC3409033.

40. Minh BQ, Nguyen MA, von Haeseler A. Ultrafast approximation for phylogenetic bootstrap. Mol Biol Evol. 2013;30(5):1188–95. Epub 2013/02/19. doi: 10.1093/molbev/mst024. PubMed PMID: 23418397; PubMed Central PMCID: PMCPMC3670741.

41. Price DL. Dirofilaria magnilarvatum n. sp. (Nematoda: Filarioidea) from Macaca irus Cuvier. I. Description of the adult filarial worms. J Parasitol. 1959;45:499–504. PubMed PMID: 14434843.

42. Taylor AE. Dirofilaria magnilarvatum Price, 1959 (Nematoda: Filarioidea) from Macaca irus Cuvier. II. Microscopical studies on the microfilariae. J Parasitol. 1959;45:505–9. PubMed PMID: 13837108.

43. Sasa M. Human filariasis: A global survey of epidemiology and control: University Park Press; 1976.

44. Wharton RH. Dirofilaria magnilarvatum Price, 1959 (Nematoda: Filarioidea) from Macaca irus Cuvier. IV. Notes on Larval Development in Mansonioides Mosquitoes. The Journal of Parasitology. 1959;45(5):513–8. doi: 10.2307/3274572.

45. Budiyanto A. Evaluation Study of Filariasis Limfatic Elimination Activities. Journal of Medical Science And clinical Research. 2019;7. doi: 10.18535/jmscr/v7i4.145.

46. Yilmaz E, Wongkamchai S, Ramunke S, Koutsovoulos GD, Blaxter ML, Poppert S, et al. High genetic diversity in the Dirofilaria repens species complex revealed by mitochondrial genomes of feline microfilaria samples from Narathiwat, Thailand. Transbound Emerg Dis. 2019;66(1):389–99. doi: 10.1111/tbed.13033. PubMed PMID: 30281949.

47. Ta-Tang TH, Luz SL, Merino FJ, de Fuentes I, López-Vélez R, Almeida TA, et al. Atypical Mansonella ozzardi Microfilariae from an Endemic Area of Brazilian Amazonia. Am J Trop Med Hyg. 2016;95(3):629–32. Epub 20160711. doi: 10.4269/ajtmh.15-0654. PubMed PMID: 27402517; PubMed Central PMCID: PMCPMC5014270.

48. Bain O, Mutafchiev Y, Junker K, Guerrero R, Martin C, Lefoulon E, et al. Review of the genus Mansonella Faust, 1929 sensu lato (Nematoda: Onchocercidae), with descriptions of a new subgenus and a new subspecies. Zootaxa. 2015;3918(2):151–93-–93.

49. Ta-Tang T-H, Crainey JL, Post RJ, Luz SL, Rubio JM. Mansonellosis: current perspectives. Research and reports in tropical medicine. 2018:9–24.

50. Mat Udin AS, Uni S, Rodrigues J, Martin C, Junker K, Agatsuma T, et al. Redescription, molecular characterisation and Wolbachia endosymbionts of Mansonella (Tupainema) dunni (Mullin & Orihel, 1972) (Spirurida: Onchocercidae) from the common treeshrew Tupaia glis Diard & Duvaucel (Mammalia: Scandentia) in Peninsular Malaysia. Curr Res Parasitol Vector Borne Dis. 2024;5:100154. Epub 20231123. doi: 10.1016/j.crpvbd.2023.100154. PubMed PMID: 38193019; PubMed Central PMCID: PMCPMC10772378.

51. Bergner JJE, Jachowski LJA. THE FILARIAL PARASITE, MACACANEMA FORMOSANA FROM THE TAIWAN MONKEY AND ITS DEVELOPMENT IN VARIOUS ARTHROPODS. 1968.

52. McNulty SN, Fischer K, Mehus JO, Vaughan JA, Tkach VV, Weil GJ, et al. Absence of Wolbachia endobacteria in Chandlerella quiscali, an avian filarial parasite. J Parasitol. 2012;98(2):382–7. Epub 2011/10/29. doi: 10.1645/GE-2879.1. PubMed PMID: 22032328; PubMed Central PMCID: PMCPMC5000782.

53. Pomari E, Voronin D, Alvarez-Martinez MJ, Arsuaga M, Bottieau E, Luzon-Garcia MP, et al. Wolbachia bacteria in Mansonella perstans isolates from patients infected in different geographical areas: a pilot study from the ESCMID Study Group for Clinical Parasitology. Parasit Vectors. 2025;18(1):97. Epub 2025/03/11. doi: 10.1186/s13071-025-06723-0. PubMed PMID: 40065479; PubMed Central PMCID: PMCPMC11895188.

